# Metabolic brain network reorganization precedes clinical conversion in Alzheimer’s disease

**DOI:** 10.64898/2026.08.04.740599

**Authors:** Christian Limberger, Guilherme Schu, Giordana Salvi de Souza, Marco A. De Bastiani, Andrei Bieger, Gabriel Colissi-Martins, Giovanna Carello-Collar, Guilherme Povala, Luiza S. Machado, Thomas H. Schlickmann, Alzheimer’s Disease Neuroimaging Initiative, Tharick A. Pascoal, Pedro Rosa-Neto, Débora G. Souza, Eduardo R. Zimmer

**Author notes:** Data used in preparation of this article were obtained from the Alzheimer’s Disease Neuroimaging Initiative (ADNI) database (adni.loni.usc.edu). As such, the investigators within the ADNI contributed to the design and implementation of ADNI and/or provided data but did not participate in analysis or writing of this report. A complete listing of ADNI investigators can be found at: http://adni.loni.usc.edu/wp-content/uploads/how_to_apply/ADNI_Acknowledgement_List.pdf.

## Abstract

**Introduction:** Brain glucose hypometabolism is a hallmark of Alzheimer’s disease (AD), yet conventional [^18^F]-fluorodeoxyglucose (FDG) positron emission tomography (PET) analyses have limited sensitivity in preclinical stages. Metabolic brain network approaches may better capture early vulnerability preceding clinical conversion.

**Methods:** Cognitively unimpaired individuals (n = 127) from the ADNI cohort with baseline FDG-PET and amyloid (A) and tau (T) status were classified as clinically stable or converters over an average longitudinal follow-up of 5.8 years. Baseline brain FDG uptake patterns were analyzed at the regional, voxel, and network levels across AT profiles. Network density was quantified globally and within functional networks. AD biomarkers and cognitive performance were also examined.

**Results:** Conventional FDG-PET SUVr analyses failed to distinguish cognitively stable individuals from clinical converters at baseline, either at the regional or voxel levels. AT(N) biomarkers and neuropsychological performance likewise did not differ significantly between groups. In contrast, clinical converters exhibited hyperconnected metabolic networks at baseline, including within the default-mode network. These effects were consistent across A−T−, A+T−, and A+T+ groups, with network density higher in clinical converters than in cognitively stable individuals. Conversely, network density among stable individuals declined with AT progression, pointing to divergent network trajectories.

**Discussion:** Metabolic network organization analysis revealed early AD-related vulnerability beyond regional hypometabolism, even before detectable amyloid positivity, and may reflect divergent trajectories of resilience and pathological propagation preceding clinical conversion. By leveraging existing FDG-PET datasets, this framework offers a valuable opportunity to identify individuals at risk of clinical progression at scale.

## 1. Background

Hypothetical models of Alzheimer’s disease (AD) progression propose that brain amyloid-β (Aβ) deposition is an early event in the pathological cascade, preceding tau accumulation, neurodegeneration (*e.g.* brain glucose hypometabolism and atrophy), and clinical symptom onset [1,2]. However, positron emission tomography (PET) imaging studies have shown a regional overlap between brain Aβ deposition and brain glucose hypometabolism in individuals with mild cognitive impairment (MCI) and AD dementia [3–5]. This spatial overlap is particularly pronounced in brain regions partially comprising the default-mode network (DMN), a group of regions characterized by high intrinsic functional connectivity and elevated metabolic demand at wakeful rest [6–9].

Indeed, evidence from both animal and human studies converges to indicate that the spatial distribution of Aβ accumulation is closely linked to regional differences in neural activity and metabolism that precede Aβ plaque formation [10–14]. These findings suggest that intrinsic patterns of brain activity and energy demand may shape regional vulnerability to Aβ deposition. In murine models, increased interstitial concentrations of soluble Aβ correlate with elevated extracellular lactate, reflecting increased neuronal activity and metabolic demand, and predict subsequent plaque deposition in the same regions [10]. Moreover, experimental evidence suggests that neuronal hyperactivity may represent a primary dysfunction in early disease stages, mediated by soluble oligomeric forms of Aβ rather than fibrillar aggregates [11]. In humans, Aβ deposition preferentially occurs in regions exhibiting higher levels of aerobic glycolysis in young adulthood [12,13]. In addition, cognitively unimpaired (CU) older adults with greater Aβ burden demonstrate hyperconnectivity on functional magnetic resonance imaging (fMRI) [14]. Collectively, these observations suggest that in early AD stages, soluble forms of Aβ may promote neuronal hyperactivity and network-level dysfunction well before the onset of clinical symptoms, potentially contributing to cognitive decline at later disease stages [15].

In this context, the connectome, comprising brain regions linked by axonal pathways or coordinated activity, may facilitate the propagation of pathological misfolded proteins or network-level dysfunction [16]. Methodological advances, including graph theory-based brain network models, have enabled a more precise characterization of connectome organization through the representation of nodes (brain regions) and edges (biological or functional connections) [16]. Thus, metabolic brain networks derived from [^18^F]-fluorodeoxyglucose (FDG)-PET, which capture inter-individual covariance in glucose metabolism [17], provide a valuable approach to assess whether network-level metabolic alterations reflect vulnerability patterns relevant to AD.

Given that soluble Aβ species are associated with metabolic demand in early AD stages in both animal models and humans, we sought to determine whether regional and network-level metabolic alterations precede clinical conversion in CU individuals. We hypothesize that early metabolic network remodelling, driven by Aβ synaptotoxicity, precedes clinical progression in AD.

## 2. Methods

### 2.1 Participants

Data used in this study were obtained from the Alzheimer’s Disease Neuroimaging Initiative (ADNI) database (http://adni.loni.usc.edu/). The ADNI is a longitudinal study with approximately 50 sites across the United States and Canada, launched in 2003, led by Principal Investigator Michael W. Weiner, MD. The main goal of ADNI is to identify how imaging and fluid biomarkers, along with neuropsychological and clinical assessments, can be used to understand, predict, and stage AD. This effort aims to identify sensitive and specific biomarkers to aid researchers and clinicians in developing new treatments and monitoring their effectiveness, while reducing the cost and duration of clinical trials. For up-to-date information, see www.adni-info.org. Participant recruitment for ADNI is approved by the Institutional Review Board of each participating site. Written informed consent was obtained from all participants at each site. All individuals undergo a standardized diagnostic assessment that yields a clinical diagnosis of either CU, MCI, or dementia due to AD, using established research criteria. The inclusion and diagnosis criteria for ADNI-1, ADNI-GO, and ADNI-2 have been described elsewhere [18,19]. We included CU individuals who had available baseline FDG-PET scans and amyloid and tau status across the AD *continuum* (*e.g.* A-T-, A+T-, or A+T+) defined by either cerebrospinal fluid (CSF) or PET imaging. Individuals were subsequently classified as cognitively stable if they maintained their diagnosis at follow-up visits (at least 1 year follow-up) or as clinical converters if they progressed to MCI or AD dementia. Participants from both groups were age-, sex-, and APOEε4-matched. In total, we included 127 individuals in the analysis.

### 2.2 Neuroimaging

PET acquisitions were conducted in accordance with the standardized protocols established by ADNI (https://adni.loni.usc.edu/help-faqs/adni-documentation/). FDG-PET images were preprocessed to a final spatial resolution of 8 mm full-width at half maximum. PET volumes were rigidly coregistered to each participant’s T1-weighted MRI using an automated procedure. The T1-weighted MRI was subsequently linearly registered and nonlinearly normalized to the MNI152 template, and the resulting transformations were applied to the PET data to resample images into standard space. FDG-PET standardized uptake value ratio (SUVr) maps were computed using the pons as the reference region. We obtained the global average metaROI SUVr from the ADNI database, comprising the bilateral angular gyri, posterior cingular, and inferior temporal gyri [20]. Mean regional FDG SUVr values were obtained for 76 regions of interest (ROIs) using the Desikan−Killiany−Tourville (DKT) atlas. For macroregional analyses, anatomically related ROIs were grouped into frontal, temporal, parietal, occipital, insular, cingulate, and limbic (subcortical) regions [21]. Aβ burden was assessed using [¹⁸F]AV-45 (florbetapir) PET and Aβ SUVr values were calculated using the cerebellar gray matter as the reference region. We obtained the Aβ global cortical composite SUVr from ADNI, with an Aβ positivity threshold set at 1.11 SUVr [22,23]. All PET scans were retrieved within 90 days of the corresponding clinical visit. Hippocampal volumes were derived directly from MRI data available in the ADNI database [24]. Further details on image acquisition and preprocessing are provided in the ADNI Procedures Manual.

### 2.3 Metabolic brain networks

#### 2.3.1 Network construction

Group metabolic brain networks were constructed by computing, across subjects, Pearson’s correlations of FDG SUVr values between all pairs of the 76 ROIs from the DKT atlas, using a multiple-sampling scheme previously stablished by our group [17]. Parameters were set as significance α = 0.05 (FDR), matrix threshold θ = 0.80, correlation coefficient threshold = 0, bootstrap samples = 2000, mean representation, and adaptive synthetic sampling approach (ADASYN) for imbalanced dataset correction [17]. These correlations reflect relationships between brain regions across subjects and are not based on canonical resting-state networks; rather, they follow a graph-theoretical approach in which the network captures interdependencies among all brain regions. Metabolic brain networks were represented as 76 × 76 adjacency matrices comprising statistically significant correlation coefficients.

#### 2.3.2 Network analyses

Graph theoretical parameters were estimated across 2000 bootstrap-derived network constructions for each group. We calculated network metrics such as density, global efficiency, and average clustering coefficient. For a detailed description of their computation, please refer to the corresponding methodological reference [17,25]. In summary, network density was calculated as the proportion of statistically significant connections relative to the total number of possible connections in the adjacency matrix and was used as a global measure of metabolic connectivity. Global efficiency quantified the efficiency of information transfer across the whole-brain network, whereas the average clustering coefficient measured the tendency of nodes to form local clusters, reflecting local segregation.

We further calculated network density within the seven Yeo functional networks [26] by restricting connections to regions belonging to each functional system and computing density scores within these subnetworks: DMN, dorsal attention (DAN), somatomotor (SMN), visual (VIS), frontoparietal (FPCN), limbic (LIM), and ventral attention (VAN) networks.

Between-group differences were quantified using the empirical distributions of network metric differences and their corresponding 95% confidence intervals (95% CI). For each bootstrap iteration, we calculated the relative percentage difference between groups (clinical converters *vs*. cognitively stable individuals), yielding empirical distributions of standardized between-group differences. Distributional characteristics were visualized using raincloud plots combining kernel density estimates and boxplots. Statistical uncertainty was summarized through empirical 95% CI derived from the bootstrap distributions, with significance defined as an interval that did not include zero.

### 2.4 Fluid biomarkers

CSF samples were obtained via lumbar puncture in accordance with ADNI protocols and analyzed using the Elecsys electrochemiluminescence immunoassay (Roche Diagnostics) to quantify Aβ_42_ and phosphorylated tau (pTau)181 levels. Established ADNI cutoffs were applied to classify biomarker status, with CSF Aβ_42_ concentrations below 977 pg/mL and CSF pTau181 concentrations above 24 pg/mL indicating biomarker positivity [27,28]. For data visualization, CSF Aβ_42_ concentrations above the measuring range (200-1700 pg/mL) were extrapolated using the raw signals and calibration curve [28]. For the analysis, Aβ_42_ values above the measuring range were set to the technical limit (16% of observations). Plasma neurofilament light chain (NfL) was analyzed by the Single Molecule array method (Simoa, Quanterix) at the Clinical Neurochemistry Laboratory, University of Gothenburg, Sweden [29].

### 2.5 Neuropsychological assessments

The neuropsychological assessments included measures of (i) global cognition: Montreal Cognitive Assessment (MOCA) [30], and Mini-Mental State Examination (MMSE) [31]; (ii) global AD cognitive performance scales: Alzheimer’s Disease Assessment Scale (ADAS) - Cognitive Subscale 11, ADAS 13, and ADAS Q4 (Word Recognition, Task 4 of the Cognitive Subscale 11 items) [32], and the ADNI-modified Preclinical Alzheimer’s Cognitive Composite (mPACC), which detects early cognitive decline by integrating performance in episodic memory (ADAS-Cog Delayed Recall and Logical Memory Delayed Recall), processing speed or set-shifting (Digit Symbol Substitution or Trail Making Test B), and global cognition (MMSE) [33]; (iii) episodic memory: Rey Auditory Verbal Learning Test (RAVLT) learning, immediate recall, forgetting, and percentage forgetting [34]. All cognitive test scores were standardized as z-scores (centered on cognitively stable individuals) and harmonized so that higher values represent better performance. Complete details of the neuropsychological battery and assessment methods can be found in the ADNI Procedures Manual.

### 2.6 Statistical analyses

Sample demographics were compared using t-tests or Wilcoxon tests for continuous variables and χ² tests for categorical variables. We performed linear models to evaluate the clinical conversion effect (converter *vs.* stable) on each outcome (FDG, AD biomarkers, or neuropsychological assessments). Models were adjusted for age, sex, and AT status (A-T-, A+T-, A+T+). Voxel-wise analyses were performed with the RMINC package (*v.1.5.3.0*) in R, using the same linear regression model approach. Multiple-comparison correction was performed using random field theory, with thresholds for T values reported in the figure legends. We generated the group-averaged FDG-PET images using the MINC Toolkit (*v.1.9.16*). Between-group differences in FDG SUVr ROI distributions were assessed using a two-sample Kolmogorov-Smirnov test.

We applied an unsupervised clustering approach to evaluate whether baseline regional FDG-PET measures could identify metabolic subgroups associated with subsequent clinical conversion [35]. Specifically, regional FDG-PET SUVR values were first standardized (z-scored) across individuals. Principal component analysis (PCA) was then used for dimensionality reduction, and the retained principal components were used as input features for k-means clustering. The number of clusters was set a priori to two, corresponding to the expected groups of cognitively stable individuals and clinical converters, without using clinical outcome during model fitting. Cluster membership was subsequently compared between cognitively stable individuals and clinical converters using Pearson’s chi-square test with Yates’ continuity correction to assess whether the identified clusters were associated with clinical outcome.

For all other analyses, a two-tailed Student’s t test was performed. We considered P < 0.05 as statistically significant. All analyses were performed in R (*v. 4.5.1)*.

## 3. Results

### 3.1 Demographic characteristics

The study included 127 participants, comprising 62 cognitively stable individuals and 65 clinical converters. Across the overall cohort and within AT profiles, stable individuals and converters were comparable in age, sex, and years of education. APOEε4 carriers were numerically more frequent among clinical converters, particularly within A+T- and A+T+ individuals, although this difference did not reach statistical significance. Most clinical converters progressed to MCI (95.4%), with an average time to conversion of approximately 5 to 7 years across AT status. The detailed demographic information is presented in **Table 1**.

**Table 1.** Demographics at baseline.

|  | Overall |  | A-T- |  | A+T- |  | A+T+ |  |
| --- | --- | --- | --- | --- | --- | --- | --- | --- |
|  | Stable | Converter | Stable | Converter | Stable | Converter | Stable | Converter |
| Individuals, no. | 62 | 65 | 28 | 28 | 16 | 16 | 18 | 21 |
| Female, no. (%) | 30 (48.4) | 30 (46.2) | 10 (35.7) | 10 (35.7) | 8 (50) | 7 (43.8) | 12 (66.6) | 13 (61.9) |
| Age, years, mean (SD) | 74.6 (5.5) | 74.8 (5.4) | 72.4 (5.3) | 72.6 (5.6) | 76.2 (4.7) | 76.3 (4.6) | 76.4 (5.3) | 76.5 (4.8) |
| Education, years, mean (SD) | 16.4 (2.8) | 15.9 (2.8) | 17 (2.6) | 15.9 (2.8) | 16.2 (3.2) | 15.8 (2.7) | 15.8 (2.8) | 16 (2.9) |
| APOEε4 carriers, no. (%) | 16 (25.8) | 23 (35.4) | 3 (10.7) | 3 (10.7) | 5 (31.3) | 9 (56.3) | 8 (44.4) | 11 (52.4) |
| Conversion to MCI, no. (%) | - | 62 (95.4) | - | 27 (96.4) | - | 16 (100) | - | 19 (90.5) |
| Conversion to AD dementia, no. (%) | - | 3 (4.6) | - | 1 (3.6) | - | 0 (0) | - | 2 (9.5) |
| Onset of conversion, years, mean (SD) | - | 5.8 (3.6) | - | 6.7 (4.1) | - | 5.3 (3.2) | - | 5 (3.1) |

Comparisons between cognitively stable individuals and clinical converters were performed using t-tests or Wilcoxon tests for continuous variables and $\chi^2$ tests for categorical variables, within each AT status and overall. No significant differences were observed between groups for any variable ( $P > 0.05$ ). Onset of conversion, and conversion to MCI/dementia are reported for clinical converters only and were not subject to between-group statistical comparison. Abbreviations: A, amyloid; MCI, mild cognitive impairment; SD, standard deviation; T, tau.

### 3.1 Metabolic brain network hyperconnectivity distinguishes clinical converters from cognitively stable individuals

Group-averaged FDG-PET brain maps were visually comparable between cognitively stable individuals and clinical converters (**Fig. 1A**). Consistent with this observation, the voxel-wise comparison did not reveal any prominent significant clusters distinguishing the two groups (T > 3.16, RFT-corrected) (**Fig. 1B**). At the regional level, no marked differences were observed, as reflected by the FDG metaROI SUVr values (P = 0.9234) (**Fig. 1C**) and the largely overlapping group distributions of SUVr values across ROIs, with a small magnitude of divergence between the two distributions (Kolmogorov-Smirnov test, D = 0.041, P < 0.001) (**Fig. 1D**). When FDG SUVr values were aggregated into clusters corresponding to cerebral macroregions, clinical converters exhibited significantly higher values in the frontal lobe compared with cognitively stable individuals (two-tailed T-test, P < 0.05; **Fig. 1E**). To capture subject-level metabolic organization, we applied unsupervised clustering to FDG profiles to try to identify distinct metabolic patterns; however, this approach did not reveal distinct clustering separating cognitively stable individuals from clinical converters (χ²_(1)_ = 0.01, P = 0.921) (**Fig. 1F**).

**Figure 1.**
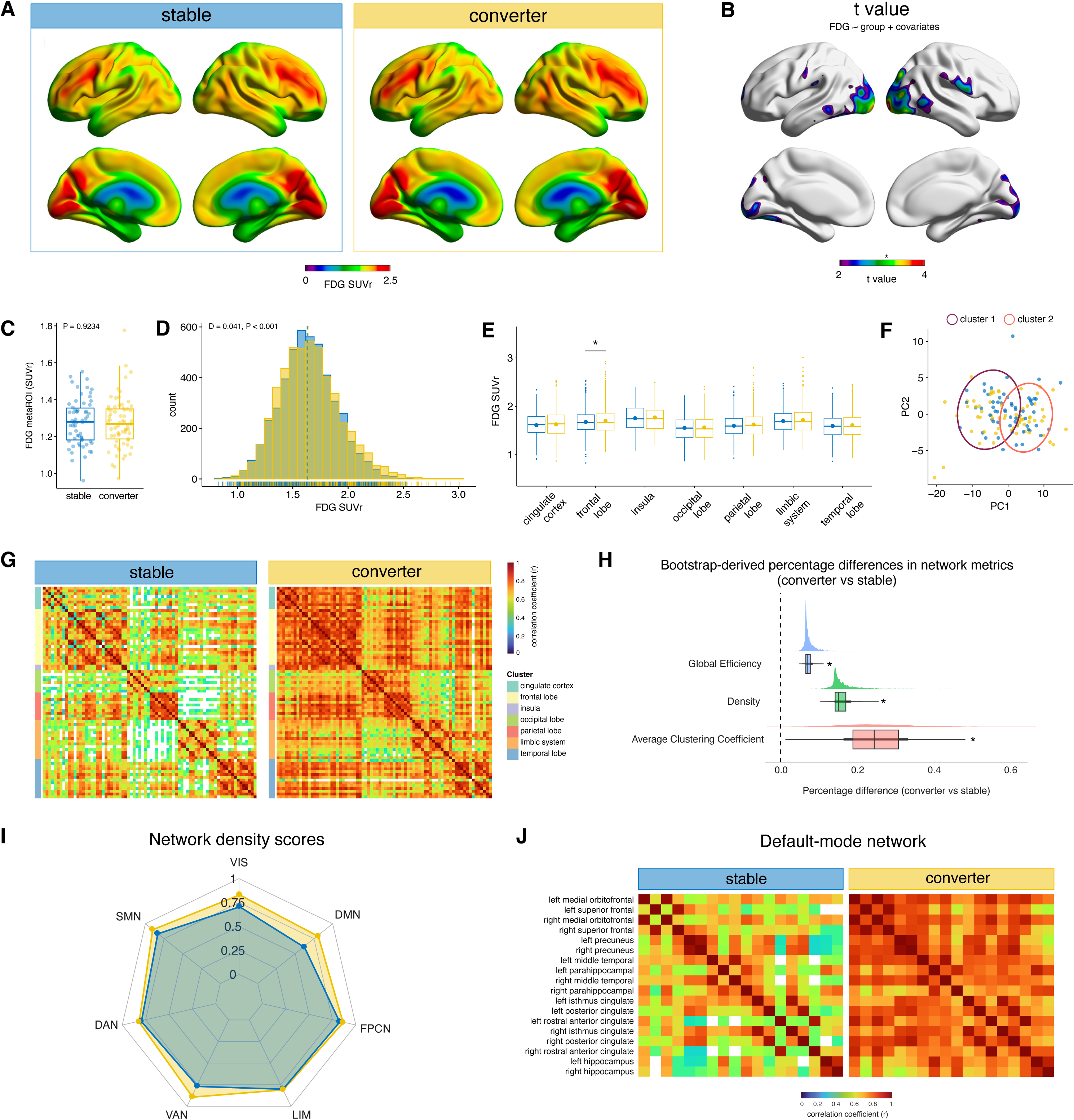
Comparison of FDG uptake and metabolic network organization between cognitively stable individuals and clinical converters. **(A)** Mean cerebral FDG uptake maps. **(B)** Voxelwise FDG-PET differences between clinical converters and cognitively stable individuals (*T > 3.16, RFT-corrected). **(C)** FDG metaROI values across groups. Boxplots indicate the median and interquartile range. **(D)** Histogram of global distributions of regional FDG SUVr in stable and converter groups, with small difference between groups (Kolmogorov-Smirnov test: D = 0.041, P < 0.001). **(E)** Mean FDG uptake by brain macroregions, with higher values in the frontal lobe in clinical converters (two-tailed T-test, *p < 0.05). **(F)** PCA of regional FDG-PET metabolic profiles with unsupervised clustering. Each point represents one participant; dot colors indicate clinical diagnosis, and circles denote cluster membership derived from the analysis. **(G)** Brain metabolic networks represented as adjacency matrices comprising statistically significant correlation coefficients. The 76 DKT brain regions are grouped by cerebral cluster, represented with different colors in the left bar. **(H)** Raincloud plots showing empirical distributions of bootstrapped percentage differences (converters vs. stable) for network metrics. Boxplots indicate the median and interquartile range; the dashed line marks no difference (0%). Asterisks denote metrics with 95% CI excluding zero (significant between-group difference). **(I)** Within-network connectivity across the seven Yeo functional networks, highlighting the largest group difference in the DMN. **(J)** Regional connectivity within the DMN, demonstrating stronger connectivity patterns in clinical converters. Abbreviations: DAN, dorsal attention network; DKT, Desikan-Killiany Tourville; DMN, default-mode network; FDG, [^18^F]-fluorodeoxyglucose; FPCN, frontoparietal network; LIM, limbic network; PCA, principal component analysis; RFT, random field theory multiple comparisons correction; SMN, somatomotor network; SUVr, standardized uptake value ratio; VAN, ventral attention network; VIS, visual network.

Metabolic brain network analysis revealed pronounced alterations in network organization. Clinical converters exhibited a more interconnected metabolic network than cognitively stable individuals (**Fig. 1G**). This observation was supported by graph-theoretical parameters quantified across multiple bootstrap sampling iterations. The central tendency of the bootstrap distributions indicated increases in clinical converters relative to cognitively stable individuals across several metrics: network density (16%; 95% CI, 12–26%), average clustering coefficient (25%; 95% CI, 9–45%), and global efficiency (7%; 95% CI, 6–11%) (**Fig. 1H**).

Analysis of connectivity within the seven Yeo functional networks indicated that the DMN exhibited the largest between-group difference in network density (stable: 0.62; converters: 0.80) (**Fig. 1I**). Subsequent examination of regional connectivity within the DMN revealed consistently stronger bilateral connectivity in individuals who subsequently converted diagnosis (**Fig. 1J**).

### 3.2 Metabolic brain network alterations precede detectable changes in baseline AT(N) biomarkers and cognition

Our analyses revealed no significant differences between cognitively stable individuals and clinical converters at baseline in amyloid (CSF Aβ_42_ and Aβ-PET), tau (CSF pTau181), and neurodegeneration biomarkers (plasma NfL and hippocampal volume) (**Fig. 2A**). Similarly, there were no significant differences between cognitively stable individuals and clinical converters at baseline in neuropsychological assessment scores, either reported as z-scores (**Fig. 2B**) or as β estimates from linear models regressing each z-scored test on clinical group, adjusted for age, sex, and AT profile (**Fig. 2C**).

**Figure 2.**
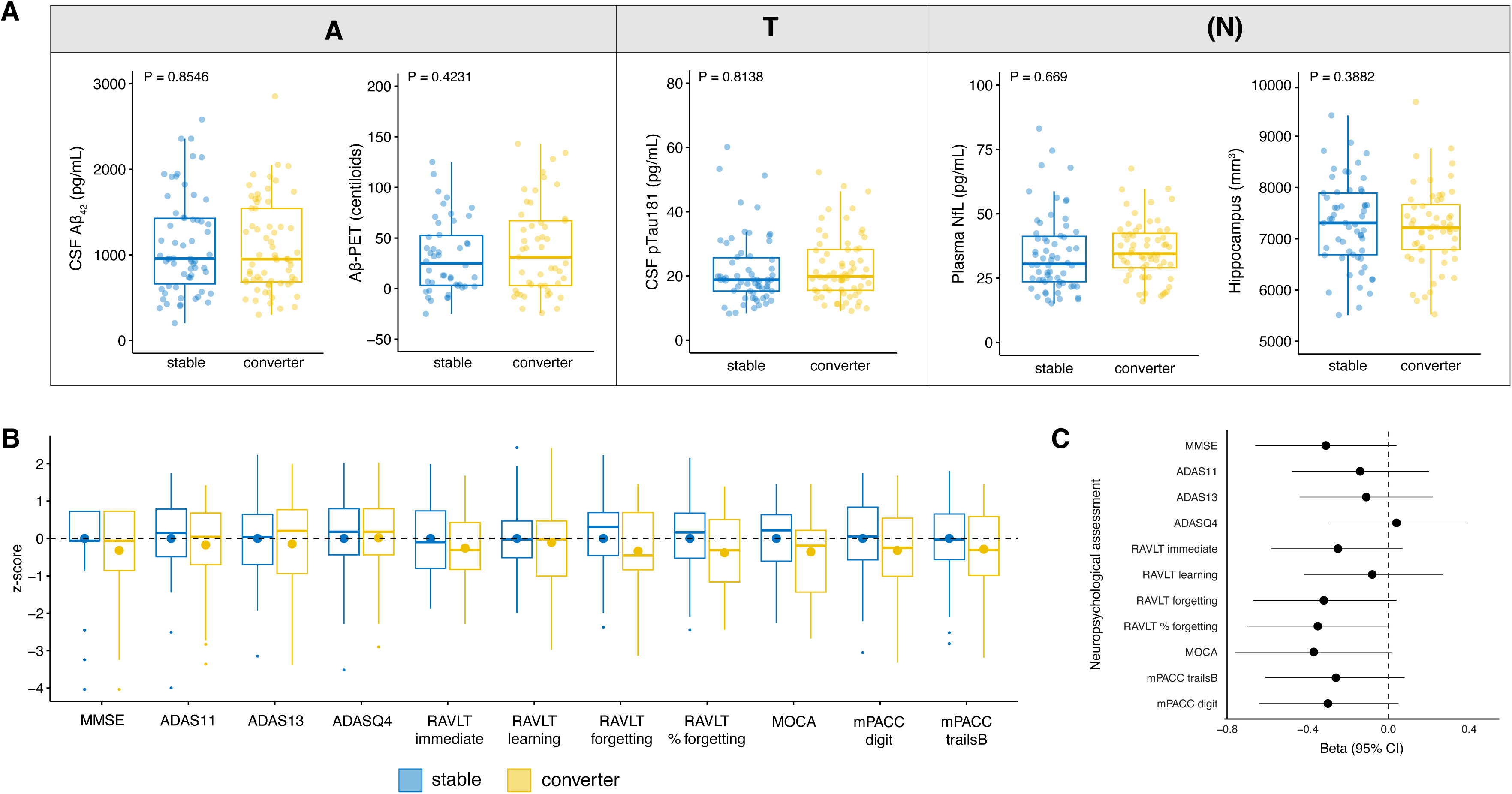
AT(N) biomarkers and neuropsychological assessments in cognitively stable individuals and clinical converters at baseline. **(A)** Levels of AT(N) biomarkers across groups (CSF Aβ_42_, Aβ-PET, CSF pTau181, plasma NfL, and hippocampal volume). Boxplots show the median and interquartile range. **(B)** Neuropsychological test scores, harmonized as z-scores centered on cognitively stable individuals, with lower values representing worse performance across multiple domains, including global cognition, executive function, and episodic memory. Boxplots show the median and interquartile range, and dots represent the mean. **(C)** β estimates from linear models (adjusted for age, sex, and AT profile) corresponding to the z-scores shown in (B). Abbreviations: A, amyloid; ADAS, Alzheimer’s Disease Assessment Scale–Cognitive Subscale; MMSE, mini-mental state examination; mPACC, ADNI-modified Preclinical Alzheimer’s Cognitive Composite; MOCA, Montreal Cognitive Assessment; N, neurodegeneration; NfL, neurofilament light chain; T, tau;

### 3.3 Amyloid and tau pathologies differentially shape metabolic brain network organization

We next stratified cognitively stable individuals and clinical converters by amyloid and tau positivity status (A-T-, A+T-, and A+T+) and evaluated their FDG-PET uptake patterns and metabolic brain network organization. When analyzing FDG brain maps, no significant clusters at the voxel-level were identified across AT profiles comparing clinical converters with cognitively stable individuals (**Fig. 3A-B**).

**Figure 3.**
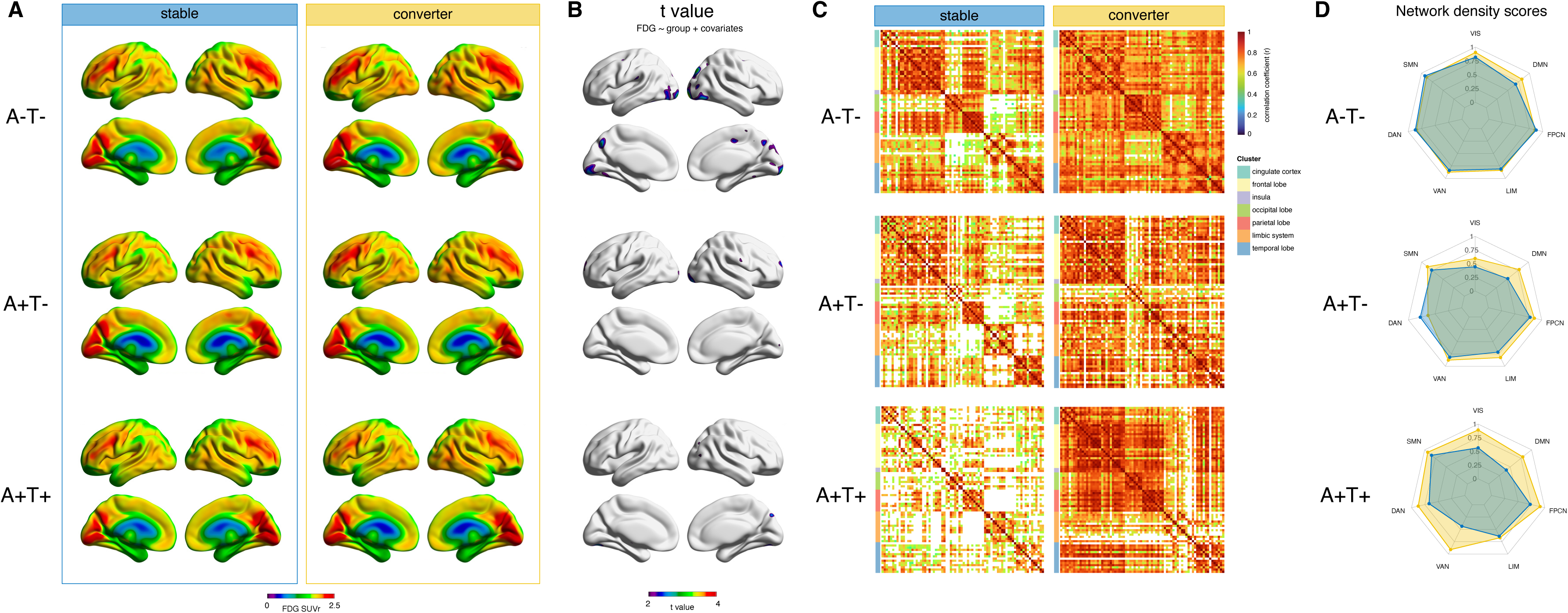
FDG maps and metabolic networks stratified by amyloid and tau positivity. **(A)** Mean cerebral FDG uptake maps stratified by amyloid (A+) and tau (T+) positivity. **(B)** Voxel-wise FDG-PET differences between clinical converters and cognitively stable individuals across AT status (RFT correction: A-T-, T > 3.25; A+T-, T > 3.40; A+T+, T > 3.34). **(C)** Brain metabolic networks of cognitively stable individuals and clinical converters. **(D)** Within-network connectivity across the seven Yeo functional networks, demonstrating increased densities in converters. Abbreviations: DAN, dorsal attention network; DMN, default-mode network; FDG, [^18^F]-fluorodeoxyglucose; FPCN, frontoparietal network; LIM, limbic network; RFT, random field theory multiple comparisons correction; SMN, somatomotor network; SUVr, standardized uptake value ratio; VAN, ventral attention network; VIS, visual network.

However, at the network level, clinical converters across all AT groups exhibited increased network density compared with cognitively stable individuals. More specifically, stable A+T− and A+T+ individuals presented hypoconnected networks relative to the other groups. Notably, A+T+ converters presented pronounced hyperconnectivity predominantly in neocortical regions, with limited involvement of limbic areas (**Fig. 3C**).

Network density scores across the seven Yeo functional networks are shown in **Fig. 3D**. In the A−T− group, metabolic network density profiles were largely comparable between cognitively stable individuals and clinical converters, although converters exhibited higher density across the DMN and VIS. By contrast, in the A+T− group, cognitively stable individuals exhibited a slight reduction in network density across nearly all functional systems relative to clinical converters. In the A+T+ group, clinical converters displayed widespread increases in network density across almost all functional networks, except for the LIM and SMN networks, which showed slightly changes compared to the cognitively stable group.

## 4. Discussion

In this study, we demonstrate that brain metabolic hyperconnectivity precedes detectable AD biomarker abnormalities and clinical diagnosis in CU individuals who later convert to cognitive impairment. Conventional ROI- and voxel-wise FDG-PET analyses failed to detect baseline differences between cognitively stable individuals and clinical converters. In contrast, network-level analyses revealed marked disruptions in metabolic connectivity, indicating that large-scale metabolic network reorganization may represent an early and sensitive marker of AD progression even before biomarker positivity.

Group-averaged FDG brain maps, regional FDG SUVr distributions, and unsupervised clustering of regional FDG uptake profiles did not distinguish clinical converters from cognitively stable individuals. The absence of group differences at the voxel and ROI levels is consistent with prior reports indicating that univariate FDG-PET measures have limited sensitivity to detect subtle metabolic changes during early and mild stages of dementia [36–38]. Parcellation of regional FDG SUVr values into brain lobes revealed subtle hypermetabolism in clinical converters, suggesting that modest increases in metabolic demand may already be present. This observation is consistent with studies proposing that FDG hypermetabolism may precede hypometabolism [39], the typical metabolic signature of AD.

In fact, regional glucose hypometabolism in AD does not necessarily reflect isolated local dysfunction but may arise from AD pathological processes affecting anatomically or functionally connected regions. More specifically, human neuroimaging studies have shown that posterior cingulate hypometabolism, a classical hallmark of prodromal AD, is associated with local and remote effects of atrophy and Aβ load, a relationship likely mediated by disruption of long-range white matter pathways such as the cingulum bundle [40–42]. Supporting evidence from tau- and Aβ-PET studies indicates that metabolic dysfunction can propagate along functional networks rather than remaining confined to regions of local pathology [43,44]. Together, these findings corroborate a concept established in both fMRI and FDG-PET connectomics, where early disease stages are characterized by disrupted inter-regional coordination despite preserved regional signals [7,45].

In contrast to the regional evaluation, metabolic brain network analyses uncovered biologically meaningful differences. Clinical converters exhibited overall topological connectivity patterns similar to those of cognitively stable individuals, but with increased density. This finding suggests that early asymptomatic disease stages may be associated with a state of network hyperconnectivity, potentially reflecting compensatory mechanisms, increased neuronal or glial activity, or altered neurovascular–metabolic coupling [46,47]. Patterns of increased metabolic connectivity have been reported in cross-sectional studies of AD, both in early and late stages of the disease. More specifically, Arnemann *et al.* constructed cross-sectional metabolic brain networks by computing direct pairwise correlations of FDG uptake and reported increases in metabolic correlations, along with network desegregation, in normal aging relative to both younger adults and individuals with AD [48]. Chie *et al.* reported that, in individuals with AD, whole-brain metabolic connectivity exhibited increased connection strength and clustering coefficients, which were interpreted as evidence of brain network reorganization reflecting compensatory responses to pathological disruption [49]. The authors interpret the findings from both studies as reflecting a compensatory reorganization of the metabolic network that may precede eventual network collapse. These findings suggests that early pathological processes may not manifest as regional abnormalities but rather as alterations in the coordination of metabolic activity across brain networks.

Clinical converters exhibited increased network density, global efficiency, and average clustering coefficient, suggesting that metabolic networks became more integrated and locally interconnected. This pattern indicates that the observed hyperconnectivity was not limited to long-range integration alone but was accompanied by a parallel increase in local clustering, consistent with a broader reorganization of network topology. Among the seven Yeo functional networks, the DMN exhibited the largest between-group differences. Progressive disruption of DMN functional connectivity, beginning even in the early stages of MCI, has been consistently reported in fMRI studies [6,50–52]. Given the DMN’s high metabolic demand, central role in cognition, and high vulnerability to Aβ [7], these results reinforce the notion that DMN metabolic connectivity may represent a link between early pathology and downstream cognitive decline.

In our cohort, however, baseline neuropsychological performance did not differ significantly between stable and converter groups. Linear models showed marginal β estimates for some measures (*e.g.*, RAVLT % forgetting, P = 0.0525), likely reflecting limited statistical power given the sample size. Nonetheless, the direction of this estimate might suggest subtle cognitive deficits in domains associated with early AD-related changes, particularly episodic memory [53]. In contrast, metabolic network analysis detected hyperconnectivity differences between groups already at baseline. Interestingly, no group differences were observed across AT(N) biomarkers in the overall cohort. Together, these findings suggest that vulnerability to AD may initially manifest at the metabolic network level, preceding detectable changes in cognitive performance, AT(N) biomarkers, or regional FDG metabolism.

Stratification by AT profiles further revealed that metabolic network alterations are modulated by underlying Aβ and tau pathology. In general, clinical converters presented higher network density than cognitively stable individuals within each AT status. Interestingly, this hyperconnectivity was also observed in the A-T- group, suggesting that metabolic network reorganization may emerge before detectable AD biomarker abnormalities. This observation aligns with emerging evidence that functional and metabolic network changes can precede classical molecular hallmarks of AD [48]. Further analyses should investigate whether Aβ accumulation propagates within regions comprising the DMN [54].

When comparing different AT profiles within clinical converters, A+T− individuals exhibited reduced network density while preserving architectural patterns similar to A−T− individuals. This network weakening may represent an intermediate stage in which Aβ plaque-related synaptic dysfunction begins to compromise large-scale metabolic integration [14]. In contrast, A+T+ converters demonstrated pronounced hyperconnectivity predominantly within neocortical systems, with relative sparing of limbic networks. This pattern may be related to the typical course of tau pathology progression, which begins in the entorhinal cortex and hippocampus (Braak stages I–II), and only later spreads to limbic regions (Braak stages III–IV) and association neocortices (Braak stages V–VI) [55]. Given that limbic involvement is commonly associated with MCI and early clinical manifestations [56,57], the partial disconnection of limbic regions while neocortical networks still exhibit hyperconnectivity may reflect compensatory mechanisms in response to region-specific vulnerability to pathological spread. Comparable coexistence of hyper- and hypo-connectivity across networks has been described in longitudinal FDG-PET studies, where early hyperconnectivity, particularly outside core limbic systems, was associated with faster progression to dementia [49,58].

In contrast, the network density in cognitively stable individuals decreased with AT progression. Such a pattern may reflect a resilience-related reconfiguration that preserves cognitive function by limiting maladaptive metabolic coupling. In contrast to clinical converters, who exhibit early network hyperconnectivity, cognitively stable individuals may downregulate inter-regional metabolic synchronization to avoid energetically costly overactivation [59]. This reduction in network density may also indicate a partial uncoupling between molecular pathology and metabolic coordination, thereby limiting network-mediated propagation of dysfunction and supporting cognitive stability. This interpretation aligns with the cascading network failure model proposed for functional connectivity, in which hyperconnectivity is thought to reflect a load-shifting process rather than beneficial compensation. In this model, the processing burden of a failing subsystem is transferred to hub regions, resulting in connectivity overload that precedes further structural and functional decline [60]. Extending this framework to the metabolic domain, the declining network density observed in cognitively stable individuals may indicate that this overload state is not being engaged, either because the initiating failure has not yet occurred or because the network is being actively reconfigured to limit the hub recruitment through which such cascading dysfunction would otherwise propagate.

Notably, this work is the first to apply a multiple-sampling scheme framework to generate stable and reproducible metabolic brain networks from reduced sample sizes, yielding networks that are robust to outliers and data imbalance [17]. This methodological advantage is particularly relevant in preclinical cohorts, where sample sizes are limited and conventional covariance approaches may yield unstable or biased network estimates, as highlighted in recent comparative evaluations of individual FDG-PET graph construction methods [61].

Our study has some limitations. First, its cross-sectional design precludes determining whether baseline network organization is associated with distinct trajectories of disease progression. Longitudinal network trajectories could not be evaluated because of the limited availability of follow-up FDG- and Aβ-PET scans in our cohort. Second, the relatively small sample size is mitigated by the multiple-sampling scheme approach, which is specifically designed to generate robust group-level representations of metabolic brain networks even in smaller samples. Finally, metabolic networks were derived at the group level rather than at the individual level, and thus potential prognostic information embedded in individual network variability could not be assessed.

## 5. Conclusion

Overall, our results suggest that alterations in brain network organization represent one of the earliest measurable signs in individuals who will later progress to the clinical phases of AD. The observed early hyperconnectivity and increased network density in clinical converters compared to cognitively stable individuals may reflect compensatory mechanisms or pathological hyperactivity. In Aβ-positive states, this pattern appears to be followed by network weakening, with hyperconnectivity re-emerging alongside tau accumulation. These findings highlight the added value of metabolic network approaches beyond conventional FDG-PET analyses for the understanding of downstream neuropathological spread in AD.

## Acknowledgements

Data collection and sharing for this project was funded by the Alzheimer’s Disease Neuroimaging Initiative (ADNI) (National Institutes of Health Grant U01 AG024904) and DOD ADNI (Department of Defense award number W81XWH-12-2-0012). ADNI is funded by the National Institute on Aging, the National Institute of Biomedical Imaging and Bioengineering, and through generous contributions from the following: AbbVie, Alzheimer’s Association; Alzheimer’s Drug Discovery Foundation; Araclon Biotech; BioClinica, Inc.; Biogen; Bristol-Myers Squibb Company; CereSpir, Inc.; Cogstate; Eisai Inc.; Elan Pharmaceuticals, Inc.; Eli Lilly and Company; EuroImmun; F. Hoffmann-La Roche Ltd and its affiliated company Genentech, Inc.; Fujirebio; GE Healthcare; IXICO Ltd.; Janssen Alzheimer Immunotherapy Research & Development, LLC.; Johnson & Johnson Pharmaceutical Research & Development LLC.; Lumosity; Lundbeck; Merck & Co., Inc.; Meso Scale Diagnostics, LLC.; NeuroRx Research; Neurotrack Technologies; Novartis Pharmaceuticals Corporation; Pfizer Inc.; Piramal Imaging; Servier; Takeda Pharmaceutical Company; and Transition Therapeutics. The Canadian Institutes of Health Research is providing funds to support ADNI clinical sites in Canada. Private sector contributions are facilitated by the Foundation for the National Institutes of Health (www.fnih.org). The grantee organization is the Northern California Institute for Research and Education, and the study is coordinated by the Alzheimer’s Therapeutic Research Institute at the University of Southern California. ADNI data are disseminated by the Laboratory for Neuro Imaging at the University of Southern California. G.P. receives financial support from the Alzheimer’s Association (24AARFD-1243899). M.A.B. receives financial support from the Alzheimer’s Association (AARFD-23-1148735). D.G.S. is supported by the Alzheimer‘s Association (AARF-D-22-928702) and by the Brazilian Initiative of Biomarkers for Neurodegenerative Diseases, Ministry of Health of Brazil (program 30420240118). L.S.M. is supported by Anna-Lisa och Bror Björnsons Stiftelse, Herbert och Krain Jacobssons Stiftelse (2025-Forskning-461), Gun och Bertil Stohnes Stiftelse (2025-034), Stiftelsen för Gamla Tjänarinnor (2025-365), Adlerbert Research Foundation, and Handlanden Hjalmar Svenssons Forskningsfond (2026-Forskningsfonden-612). T.A.P. is supported by National Institute on Aging (R01AG075336, R01AG073267). E.R.Z. is supported by grants from Conselho Nacional de Desenvolvimento Científico e Tecnológico (CNPq) [312410/2018-2; 435642/2018-9; 312306/2021-0; 409066/2022-2; 447074/2023-7; 409595/2023-3; 444880/2024-0], Fundação de Amparo à Pesquisa do Rio Grande do Sul (FAPERGS) [ARD 21/2551-0000673-0], Alzheimer’s Association [AARGD-21-850670], Programa de Apoio a Núcleos de Excelência (PRONEX, FAPERGS/CNPq) [16/2551-0000475-7], Instituto Nacional de Ciência e Tecnologia em Neuroproteção [408446/2024-2], Instituto Nacional de Ciência e Tecnologia em Saúde Cerebral [406020/2022-1], Instituto Serrapilheira [Serra-1912-31365; R-2401-47242], Alzheimer’s Association and National Academy of Neuropsychology [ALZ-NAN-22-928381], Secretary of Health of the State of Rio Grande do Sul [SES/RS 23/2000-0091938-0], and the Brazilian Ministry of Health [00030420240118-003490]. This study was financed in part by the Coordenação de Aperfeiçoamento de Pessoal de Nível Superior - Brasil (CAPES) - Finance Code 001.

## Conflict of interest statement

E.R.Z. has served on the scientific advisory board, as a consultant or speaker for Nintx, Novo Nordisk, Biogen, Lilly, Magdalena Biosciences, and masima. E.R.Z. and M.A.B. are co-founders and minority shareholders at masima. T.A.P. serves on a data safety monitoring board for NIH award 5 R01 AG084834 based at the University of Cincinnati. He also has served on a scientific advisory board at Johnson & Johnson. P.R.-N. provides consultancy services for Roche, Cerveau Radiopharmaceuticals, Lilly, Eisai, Pfizer, and Novo Nordisk, and also serves as a clinical trials investigator for Biogen, Novo Nordisk, and Biogen. The other authors declare that they have no competing interests.

